# Phenological shifts drive biodiversity loss in plant–pollinator networks

**DOI:** 10.1101/2020.04.03.023457

**Authors:** Mauricio Franco-Cisterna, Rodrigo Ramos-Jiliberto, Pablo Moisset de Espanés, Diego P. Vázquez

## Abstract

Plant–pollinator interactions are key for ecosystem maintenance and world crop production, and their occurrence depends on the synchronization of life-cycle events among interacting species. Phenological shifts observed for plant and pollinator species increase the risk of phenological mismatches, threatening community stability. However, the magnitudes and directions of phenological shifts present a high variability, both among communities and among species of the same community. Community–wide consequences of these different responses have not been explored. Additionally, variability in phenological and topological traits of species can affect their persistence probability under phenological changes. We explored the consequences of several scenarios of plant–pollinator phenological mismatches for community stability. We also assessed whether species attributes can predict species persistence under phenological mismatch. To this end, we used a dynamic model for plant–pollinator networks. The model incorporates active and latent life-cycle states of species and phenological dynamics regulating life-cycle transitions. Interaction structure and species phenologies were extracted from eight empirical plant–pollinator networks sampled at three locations during different periods. We found that for all networks and all scenarios, species persistence decreased with increasing magnitude of the phenological shift, for both advancements and delays in flowering phenologies. Changes in persistence depended on the scenario and the network being tested. However, all networks exhibited the lowest species persistence when the mean of the expected shift was equivalent to its standard deviation and this shift was greater than two weeks. Conversely, the highest species persistences occurred when earlier-flowering plants exhibited stronger shifts. Phenophase duration was the most important attribute as a driver of plant persistence. For pollinator persistence, species degree was the most important attribute, followed by phenophase duration. Our findings highlight the importance of phenologies on the stability and robustness of mutualistic networks.

**Author summary:** Plant-pollinator interactions involve a great number of species and are essential for the functioning of natural and agricultural systems. These interactions are facing a great number of threats. In both plants and pollinators, life-cycle events including flowering and adult emergence are triggered by environmental cues such as temperature and snowmelt. Climate change has the potential to alter the timing of these events. These *phenological shifts* generate mismatches in the timing of interacting species. Thus, plants and their pollinators may not match in time and/or space, leaving flowers unpollinated and disrupting pollinator feeding. Given that natural communities are composed of multiple species interacting in complex ways, experimentally assessing the effects of this kind of perturbation is difficult. To tackle this challenge, we simulated different scenarios of phenological shifts for several empirical communities. Our results indicate that strong shifts in the timing of life-cycle events may represent a greater risk of community collapse. Likewise, plants with short blooming periods and pollinators with short activity periods or high specialization face a greater risk of extinction.

## Introduction

Interactions between plants and their pollinators are a fundamental component of production and biodiversity in terrestrial ecosystems. The majority of angiosperm plants (87.5%) depend of animal pollination for reproduction [1], and thousands of animal species obtain food and other resources produced by flowers [2]. On the other hand, pollination plays an important role in world crop production [3]. Plant–pollinator interactions can occur only if flowers and active pollinators (e.g. adult insects) overlap in space and time, and cannot be realized if any of the mutualists is either absent or present only in a latent state such as seeds or larvae. Thus, an adequate synchronization between the life cycles of plants and pollinators is critical for the realization of mutualistic interactions.

Long–term studies indicate that the phenologies of many organisms are shifting [4–7]. One of most frequently observed phenomena has been the shifts in the occurrence of plant flowering [8], possibly due to changes in average temperatures [8, 9]. Because plant and animal phenologies are likely to respond to different climatic cues [8, 10, 11], phenological mismatches are increasingly likely [8, 12, 13], with uncertain effects on the strength and maintenance of mutualistic interactions, species abundance and persistence and ecosystem functioning [13, 14]. In spite of our increasing understanding of the population–level consequences of plant–pollinator mismatches, it is largely unknown how complex ecological communities will respond to such phenological mismatches. Furthermore, the magnitudes and directions of phenological shifts has been shown to vary both among different communities and among species of the same community [9]. Thus, there is a dearth of studies exploring the community-wide consequences of plant–pollinator phenological mismatches.

The dynamics of multispecies networks experiencing phenological shifts involve broad temporal scales, broad enough to make empirical research unfeasible. Thus, to study these networks using mathematical models is a reasonable alternative. Appropriate models should represent the long–term as well as the phenological dynamics of multiple interacting species [15, 16]. In addition, life-cycle transitions play a major role in phenological dynamics and we see as necessary to represent them in the model. Yet, most of the modeling efforts conducted so far to understand the consequences of phenological shifts on community dynamics have excluded these desirable properties. For example, some studies [17, 18] have considered only single species systems. In [19] they relied on a static analysis to study multispecies networks. In [20, 21], the adopted approach is dynamical but it considers a single pair of interacting species. None of them consider life-cycle transitions.

Here we assess the consequences of plant–pollinator phenological mismatches for community stability using mathematical modeling. To this end, we built a dynamic model for plant–pollinator networks that incorporates multiple life-cycle states (active and latent) of species and phenological dynamics regulating life-cycle transitions. This model is then used to assess numerically the effects exerted by several scenarios of phenological mismatch between flowering plants and their pollinators on the long-term stability of mutualistic networks. Our *in silico* experiments were parameterized based on empirical plant–pollinator networks with known structure of interactions and phenologies of their constituent species. Using this modeling approach, we tested the hypothesis that increasing plant–pollinator mismatches would result in decreased persistence of interacting species. Furthermore, we also assessed whether relevant species and interaction attributes (species degree, phenological span, etc.) could predict species-specific persistence probability under phenological mismatch.

## Materials and methods

### Empirical networks

For conducting this study we used eight mutualistic plant–pollinator networks. Each network corresponds to a community at a given location and during a sampling period. There were three locations as shown in Table 1. Empirical data consist of: a) topology (pattern of plant–pollinator interactions), b) flowering phenophases (dates of beginning and end of flowering season of each plant species), and c) pollinator phenophases (periods of activity for each pollinator species).

**Table 1.** Database of empirical networks and phenologies used in the simulations.

| Location | Sampling date | Network size | References |
| --- | --- | --- | --- |
| Zackenberg, Greenland | Spring 1996 | 92 species: 31 plants, 61 pollinators | [22] |
| Zackenberg, Greenland | Spring 1997 | 95 species: 31 plants, 64 pollinators | [22] |
| Nahuel Huapi, Argentina | Summer 1999–2000 | 163 species: 17 plants, 146 pollinators | [23, 24] |
| Villavicencio, Argentina | Spring 2007 | 105 species: 35 plants, 70 pollinators | [25, 26] |
| Villavicencio, Argentina | Spring 2008 | 147 species: 44 plants, 103 pollinators | [25, 26] |
| Villavicencio, Argentina | Spring 2009 | 140 species: 37 plants, 103 pollinators | [25, 26] |
| Villavicencio, Argentina | Spring 2010 | 101 species: 36 plants, 65 pollinators | [25, 26] |
| Villavicencio, Argentina | Spring 2011 | 159 species: 46 plants, 113 pollinators | [25, 26] |

### Population dynamics

We use a dynamic model as a set of coupled ODEs. This system is a particular case of a more general integro–differential model published in [27]. Each plant species is described by five state–variables: adult reproductive plants (*P*), immature seeds (*I*), mature seeds (*S*), flowers (*F*) and floral resources (*N*). For simplicity, pollinators are assumed to be composed only by insect species, and are described by three state–variables: active adult pollinators (*A*), immature larvae (*E*) and mature larvae (*L*). The transitions between the state–variables are depicted in Fig 1.

**Fig 1.**
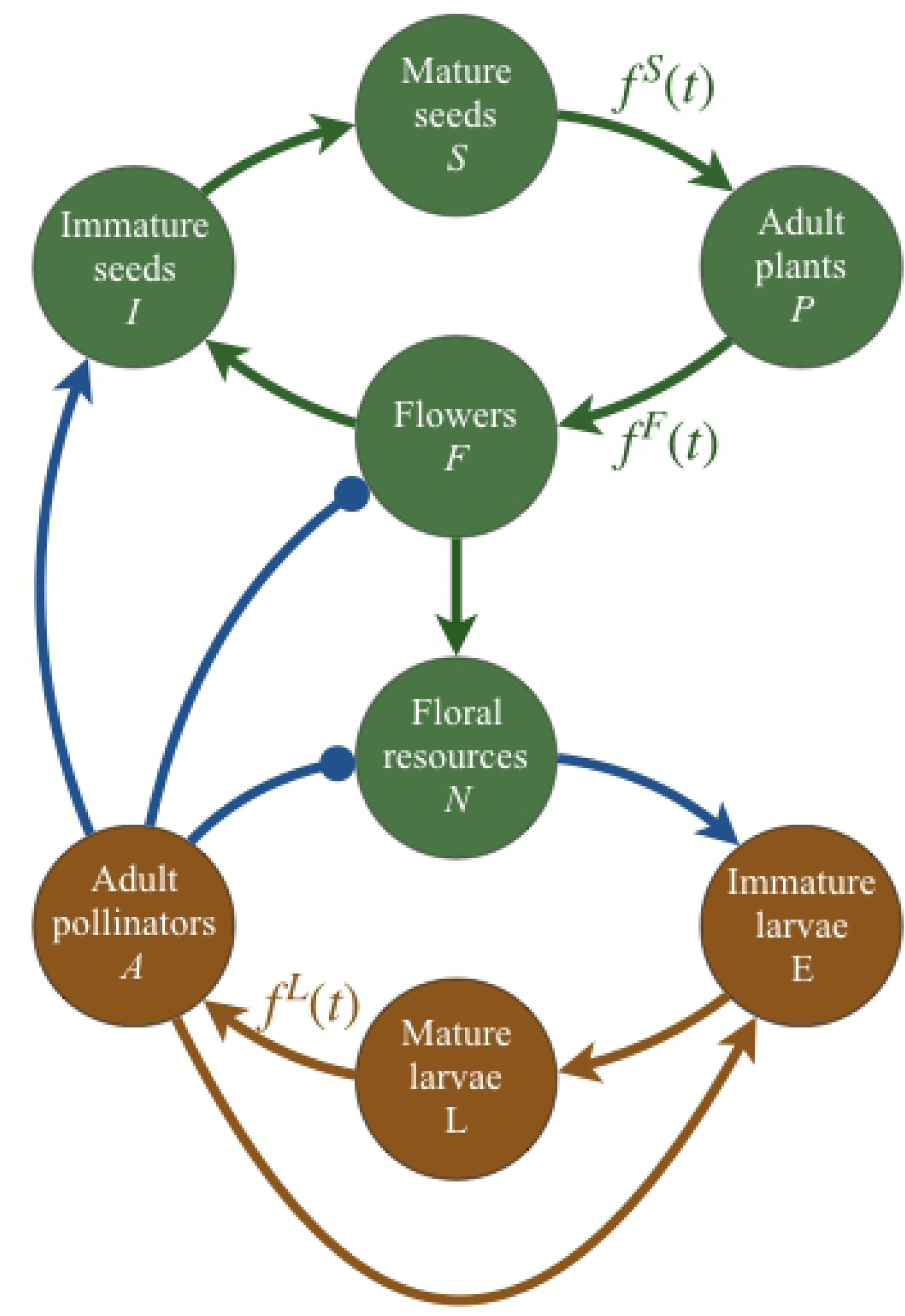
Graphical representation of the basic structure of our dynamic model. Lines ended in arrows/circles indicate a positive/negative effect of one variable on another. Green/red lines indicate life-cycle transitions within plant/pollinator species. Blue lines represent interactions between plants and their visiting pollinators. Self-effects not shown for simplicity. Those transitions marked with *f* (*t*) are governed by time–dependent functions. In these functions, superscripts *S, F* and *L* indicate seed germination, flowering and pollinator recruitment from larvae.

All state–variables are measured in biomass density, and their values vary over chronological time (*t*). Mature seeds and larvae transit to adult states when they match the appropriate subseason where environmental conditions are favorable for germination/recruitment. In our model, adult plants can produce flowers in certain period of the year, and flowers produce floral resources. Adult pollinators visit flowers and consume floral resources. After visitation, fertilized flowers are not anymore available to pollinators and immature seeds and larvae are produced (Fig. 1). Environmental favorability for germination, flowering and pollinator recruitment are described by time–dependent functions named here as “phenology functions”, *f*(*t*), which govern timing and intensity of occurrence of phenological events. Temporal dynamics of immature seed biomass is given by:

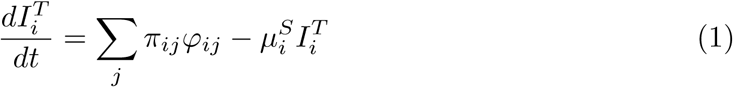

Seed production is proportional to visitation rate (*φ*) of flowers with biomass 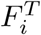 by pollinators with biomass 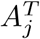. The second term represents immature seed mortality. Mature seed dynamics is given by

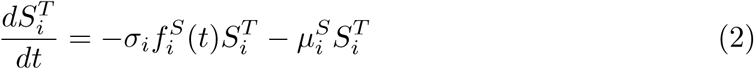

Germination phenology is governed by function 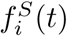 and was located in the same temporal position for all plant species. The beginning of germination was fixed to 6 weeks after the end of flowering of the latest species. The temporal duration of the germination period was set to 8 weeks for all plant species. The second term represents mature seed mortality. Biomass density growth rate of adult plants is given by:

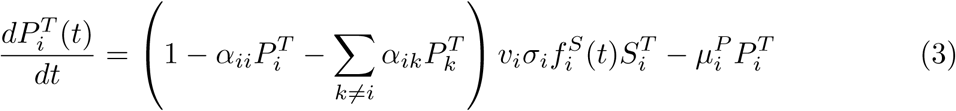

where plant biomass production due to seed germination is limited by intra- and inter-specific competition for space (term within parenthesis). The last term is plant mortality rate. Dynamics of flower biomass density is given by:

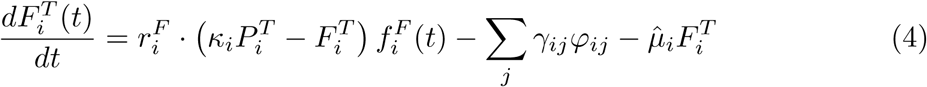

with mortality rate 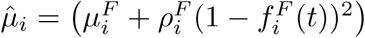, composed by basal mortality 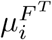 and added seasonal mortality related to *f* (*t*). This added mortality impedes flowers to persist beyond the flowering season. The first term of the equation represents the increase in flower biomass, limited by plant biomass. The second term is removal of fertile flowers due to fertilization. The third term is mortality rate, explained above. Biomass dynamics of floral resources are:

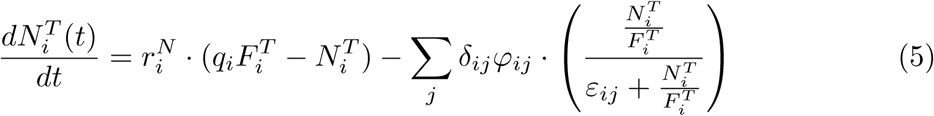

where the first term represents the increase in floral resources, which is limited by flower biomass. The second term represents resource consumption by pollinators. Resource consumption is determined by visitation rate *φ* and the amount of resources extracted by pollinators in each visit. Resource extraction per visit is a saturating function of resources per unit flower. Biomass growth rate of immature larval pollinators is given by the equation:

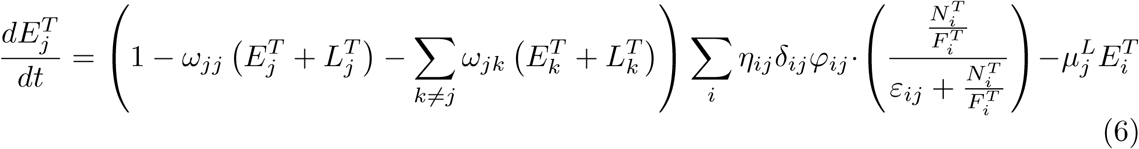

that has the same structure that equation (1) for seeds. Mature larvae follow the equation

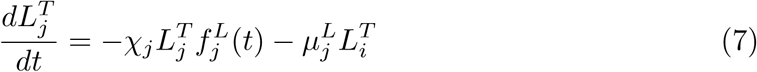

Biomass density dynamics of adults insects is governed by:

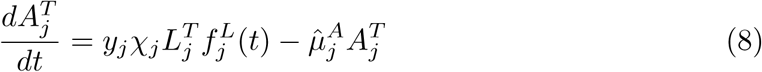

with 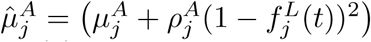, which is similar to the equation for plant biomass density (3), but without competition for space, and presents basal and seasonal mortality like flowers (4). Visitation rate of flowers by pollinators was modeled as a Beddington–DeAngelis-like functional response:

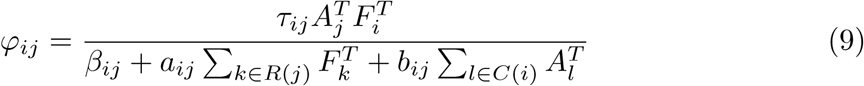

where *R*(*j*) is the set of plant species visited by pollinator *j* and *C*(*i*) is the set of pollinator species that visit plant *i*. Finally, the between-years dynamics of seeds is governed by

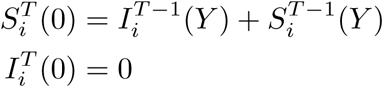

where *Y* is the length of the year. Equivalently, inter-annual dynamics of larvae is

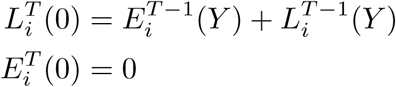

Parameter values were drawn from a random uniform distribution, centered at values taken from assessments obtained from available studies on some species, and defined and 1.25 of this value as boundaries of distribution.

### Simulation

We obtained numerical solutions for the equations using the *ode23* function of Matlab R2018b. We run every instance for 5000 weeks as a transient phase and additional 5000 weeks for the experimental phase, where treatments are applied. Initial values of biomass densities were obtained randomly from uniform distributions, centered at specific values for each state–variable (See S1 Table). State–variables were forced to zero whenever their value decreased below its given extinction threshold (See S1 Table). At the same time, a species was considered extinct when all their states–variables fell down to zero. For each replicate, the parameter values were randomly drawn from uniform distributions, centered in values taken from biologically plausible estimates or based on available literature. These values are shown in S2 Table. For each replicate, a transient simulation was run until the system reached an asymptotic oscillatory behavior, which can be considered analogous to the steady behavior in classical autonomous dynamic models. The number of species persisting at the end of this transient dynamics was recorded and used as the initial species richness for the post-transient phase.

### Experimental design

We evaluated the effects of temporal mismatch between flowering and pollinator activity on the long–term dynamics of plant–pollinator communities. To this end, we conducted three experiments. In all cases, the center of the flowering period of each species was shifted an amount of time that was randomly drawn from a normal distribution, which we call *TSD* (short for “temporal shift distribution”).

#### Experiment 1

In the first experiment, for each simulation in each year we choose an amount of temporal shift in flowering of each plant species. We conducted three variants of this experimental setting: 1. Shifting the mean of *TSD* by 25 equally–spaced levels, from −6 to 6 weeks, with standard deviation of *TSD* set to 1. Positive and negative shifts represent delayed and precocious flowering respectively. This experiment variant allows evaluating the effects produced by temporal mismatches between flowering and pollinator activity. These treatments, termed *TSD–m*, emulate phenological shifts observed in plants by [9]. 2. Increasing the standard deviation by 25 equally–spaced levels, from −6 to 6 weeks, with mean of *TSD* set to 0. This allows evaluating the effects attributable to changes in the variability of phenological shifts among species, even if the community as a whole tends to keep the central position of their phenologies. These treatments, termed *TSD–sd*, emulate phenological shifts observed in plants by [9]. 3. Shifting the mean by 25 equally–spaced levels, from −6 to 6 weeks, with standard deviation of *TSD* equal to the mean. Here, larger shifts in the mean are associated to larger increases in variability. These treatments, termed *TSD–msd*, follow the approach used in previous studies on phenological mismatches between plants and pollinators [19, 28].

Each year and for each species, we drew a new random value of phenological shift from the same *TSD*. Each treatment was replicated 50 times for each empirical network, so we run a total of 8(networks) × 3(experiment variants) × 25(levels) × 50(replicates) = 30,000 post-transient simulations. At the end of each simulation we recorded the species persistence in the community, measured as the fraction of species with positive biomass at the end of the simulation, relative to the initial set of species (after the transient phase). Species persistence was recorded for the whole set of species as well as for plants and pollinators separately. In this and subsequent experiments, we performed a non-metric multi-dimensional scaling (NMDS) analysis to evaluate differences in our results among networks. We also tested, with a Mantel test, the null hypothesis that the community persistence differ between networks from different locations but do not differ between networks sampled at different years in the same location.

#### Experiment 2

In this experiment, we consider the issue of phenological shift magnitude changing gradually through time [29]. For addressing this issue, experiment 2 differed from experiment 1 in that the expected times of flowering seasons are altered each year by an increasing magnitude. More precisely, the center of the flowering season of each species was shifted an amount of time randomly drawn from *TSD*, with standard deviation equal to the mean. The value of the mean of *TSD* was linearly increased in successive years. Maximum value of mean ranged from −6 to 6 weeks, with 25 equally–spaced levels. For this experiment we run a total of 8(networks) × 25(levels) × 50(replicates) = 10,000 post-transient simulations. At the end of each simulation we recorded species persistence in the community. These treatments were termed *TSD–msdy*.

#### Experiment 3

This experiment was designed for addressing whether plant species with earlier flowering phenophases tend to exhibit stronger phenological shifts as a response to climate change [30, 31].

These simulations are similar to those of previous experiments, but differed in that the flowering period of each plant species was shifted by an amount of time randomly drawn from a *TSD*, with standard deviation equal to the mean. In this experiment, the values of the mean of *TSD* were assigned for each species according to the starting date of its flowering. To the earliest–flowering plant we assigned the mean of *TSD* a value *M*. To the latest–flowering species we assigned zero. For all other species we used a linear interpolation. Values of *M* were set to −6, −3, 3 and 6 weeks. Each experiment was replicated 50 times for each empirical network, so we run a total of 8(networks) × 4(levels) × 50(replicates) = 1,600 post-transient simulations. At the end of each simulation we recorded species persistence in the community. These treatments were termed *TSD–msdf*.

### Predictors of species persistence

We evaluated the role of some topological and phenological attributes of the species as potential determinants of extinction risk under the analyzed scenarios of phenological shifts. Topological attributes were species’ degree, mean neighbours’ degree and variance of neighbours’ degree. Phenological attributes were phenophase duration (flowering for plants, activity period for pollinators) and mean and variance of neighbours’ phenophase duration.

For each replicate we recorded the values of species attributes to relate these values to its final status (persistent/extinct). These relations were evaluated using Random Forest (RF) models [32]. We built RFs by fitting 200 decision tree classifiers and considering all species attributes into each tree–building process. We excluded from analysis all cases where either all species went extinct or all species persisted, since no inference is possible. To assess relative importance of each attribute as a predictor of species persistence, we used the *permutation importance* metric. This metric is recommended in cases were there variations in their scale of measurement or there exists an unequal number of levels per variable [33, 34]. For each treatment and separated by plants and animals, we added the importance values of each attribute over all levels of phenological shift. We conducted the RF analysis using *scikit–learn* V.0.19.1 [35] and *rfpimp* V.1.3.3 [33] through their Python interfaces. To evaluate differences in our results among networks, we performed a NMDS analysis. We also tested, with a Mantel test, the null hypothesis that the importance values of plant and pollinator attributes differ between networks from different locations but do not differ between networks of different years in the same location.

## Results

We organize the results into two parts. First we show species persistence at the community level resulting from shifting the flowering phenology under a variety of *TSDs*. We then identify the species attributes that best explain the persistence of species subjected to phenological shift.

### Species persistence under phenological shift

The Mantel test showed that species persistences obtained for different networks sampled at diffferent times at the same location cannot be considered equivalent (*r* = 0.1494; *P* = 0.279; see S1 Fig). For this reason, we show persistence curves separated by network (Fig. 2).

**Fig 2.**
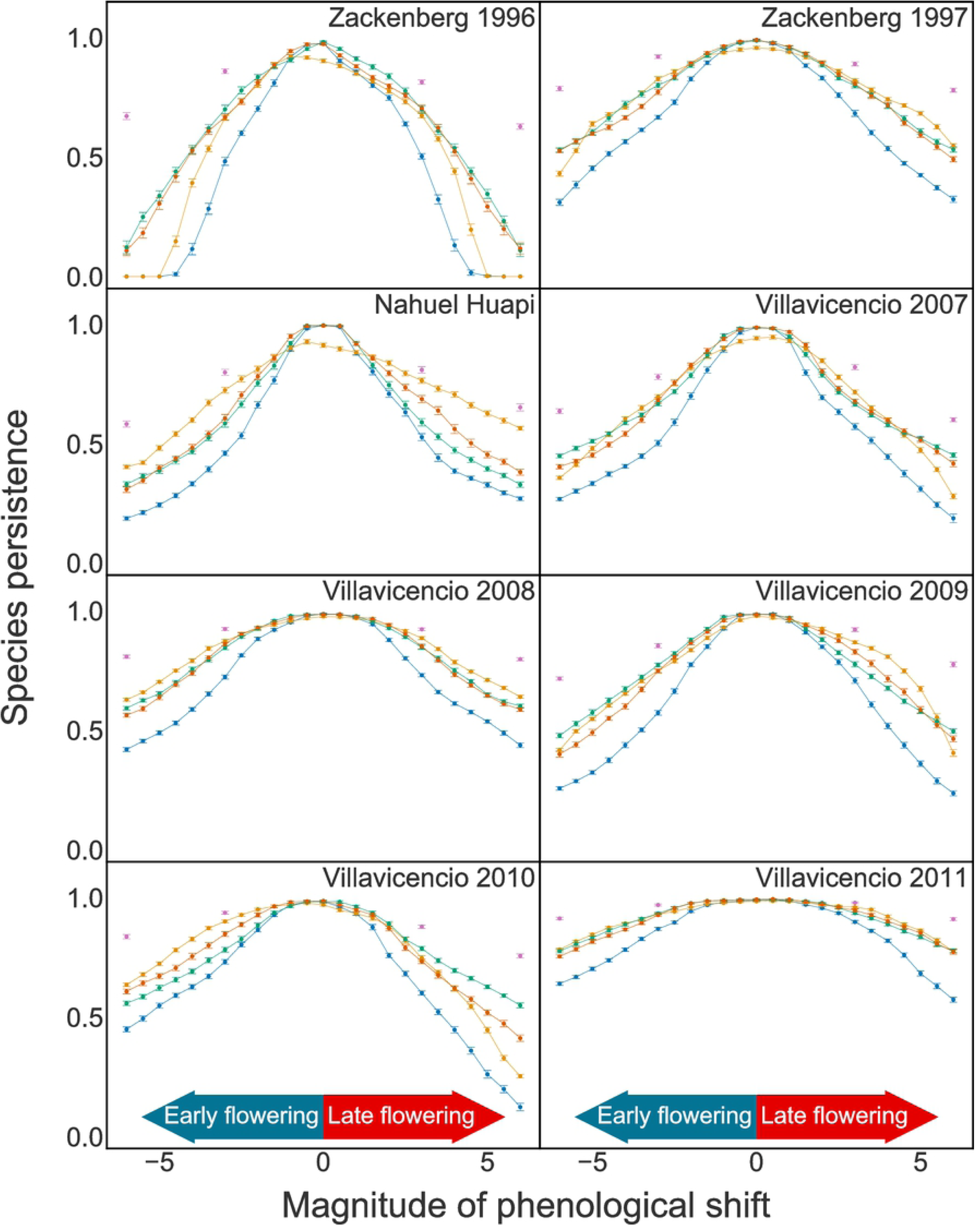
Long–term species persistence in the plant–pollinator networks under different modes and magnitudes of phenological shifts. The mode of phenological shift is indicated by colors. *TSD–msd* (blue), *TSD–m* (orange), *TSD–sd* (green), *TSD–msdy* (red), *TSD–msdf* (purple). See text for details.

For all networks and all choices of *TSD*, species persistence decreased with increasing magnitude of the phenological shift, for both advancements and delays in flowering phenologies (Fig. 2). However, in most cases, changes in persistence did not depend only on the magnitudes of phenological shifts but also on their direction: phenological advances and delays exerted different effects. Changes in persistence were also dependent on the choice of *TSD* and the network being tested. However, for all networks, the differences among the tested *TSD*s grew with increasing magnitude of the shift. For shifts of less than two weeks, *TSD–m* rendered a significantly lower species persistence for networks Zackenberg–1996, Zackenberg–1997, Nahuel Huapi and Villavicencio–2007. The other networks exhibited no differences in species persistences among the tested *TSD*s. For greater shifts (two or more weeks), for all networks, the lowest species persistences were obtained for *TSD–msd*, while the highest species persistences were obtained for *TSD–msdf*. For the other *TSD*s, their relative effects depended on both the study network and the magnitude of the shift. However, there is a discernible trend: higher persistences were usually obtained for *TSD–m*, and in fewer cases for *TSD–sd* or *TSD–msdy*.

Analyzing species persistence separately for plants and pollinators offers additional insights: while pollinator response followed the above pattern for community persistence, for plant response curves were flatter and on average higher than for pollinators, with the exception of Zackenberg–96 (see figures in S1 Appendix). Plants were less sensitive than pollinators to all *TSD*s, except for *TSD–msdf*, which drove relatively similar responses for plants and pollinators. For plants, *TSD–m* and *TSD–msd* triggered similar responses. Conversely, for pollinators, these curves fell far apart. The smallest plant persistences were found for *TSD–msd* and *TSD–m*. These results were consistent across all networks studied and independent of the shift direction.

### Explanatory attributes for persistence

The Mantel test showed that the importance values of attributes were more similar among networks from the same location than among networks from different locations (*r* = 0.5205; *P* = 0.017; see S2 Fig). Importance values indicate how well the traits predict species persistence. For this reason, we averaged the importance values among networks of the same location (Fig. 3).

**Fig 3.**
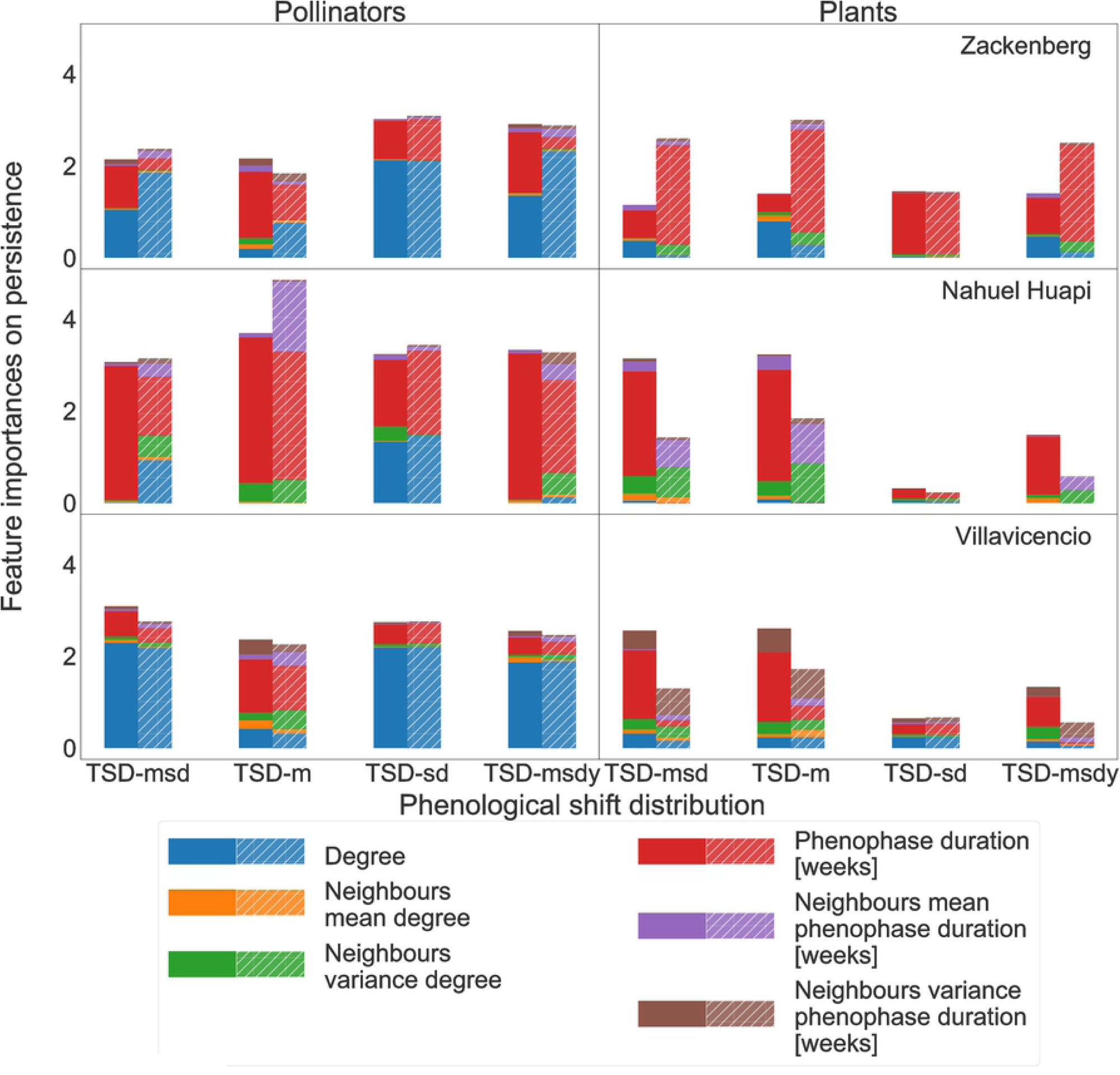
Importance measures of species attributes for the long–term persistence of pollinators and plants. Results are presented for the different locations. The categories in the x–axis represent different phenological change scenarios, modeled by different temporal shifts distribution, *TSD* (see Methods): *TSD–msd* (B), *TSD–m* (M), *TSD–sd* (V), *TSD–msdy* (Y). Solid bars: earlier flowering; dashed bars: later flowering.

Phenophase duration was the most important attribute as a driver of plant species persistence, while species degree was the most important attribute for pollinators, followed by phenophase duration (Fig. 3). The only exception to this pattern was the guild of pollinators in the Nahuel Huapi network. In this network, species persistence was largely explained by phenophase duration. Our results suggest that intrinsic species attributes are more important than those of their interaction partners as predictors of persistence. For pollinators, the direction of phenological shift and the assumed *TSD* exerted little influence on the distribution of attribute importances within treatments. For plants, the phenological shift direction affected both the magnitude of the importance of dominant attributes and the ranking of attribute importance. The first case is best seen in Zackenberg while the second one is more apparent in Nahuel Huapi and Villavicencio (Fig. 3).

The magnitude of the phenological shift had little influence in determining which attribute was the dominant one (see figures in S2 Appendix). The six exceptions among the 48 treatments were Nahuel Huapi *TSD–m* and *TSD–sd* for pollinators; Villavicencio–2009 *TSD–m* for plants; Villavicencio–2010 *TSD–m* for pollinators and plants; Villavicencio–2011 *TSD–m* for pollinators.

## Discussion

In this study we used a dynamic multispecies model that included seasonally forced biological processes to study the network consequences of phenological mismatches between pollinators an their host plants. Our results showed that phenological advancements as well as delays in plant flowering phenology lead to decreases in plant and pollinator species persistence under different modes of phenological shifts. This result is far from trivial, as it suggests that the recorded empirical set of phenophase timing of interacting species is already at some community optimum. This temporal arrangement could be attributable to co-evolutionary processes. However, this optimum could involve either plants appearing before their pollinators or vice-versa.

It may seem intuitive that this optimal arrangement involves perfect temporal matching among interacting species. However, this is not necessarily the case. In principle the optimal could involve either plants appearing before their pollinators or vice-versa. In the match/mismatch hypothesis [36], it is proposed that a resource population appearing earlier in the season favors the success of their consumers. Extrapolating to our study, this suggests that at the optimal temporal arrangement, host plants should present earlier flowering phenophases than their pollinators. By contrast, Fagan *et al.* [21] found that a higher plant population growth is obtained when pollinators recruit shortly before plant flowering. Likewise, [37] published a similar result for trophic networks. Unfortunately, their approaches considered only two and three species respectively and therefore they are not directly applicable to our networks. The first reason lies on the difficulty in quantifying basal phenophase shifts (i.e. those occurring naturally in the field before any data manipulation) among multiple interacting species. The second one is that our raw data consists of interaction records and not of independent plant and pollinator phenophases.

Of the five modes of phenological shifts, *TSD–msd* caused the largest number of extinctions. This scenario assumes that that mean and the standard deviation of flowering time change proportionally. The other shift modes tested here were less adverse for species persistence. In particular, under the realistic scenario in which earlier flowering plants are more shifted than later flowering ones, the networks showed to be relatively less sensitive to these disruptions, which is largely attributable to the higher persistence of pollinators found for this scenario. Note that our most adverse scenario *TSD–msd* was the only one used in [28] and [19], which opens the question about the outcomes of their approaches under the rest of scenarios studied here.

The decrease in species persistence caused by phenological shifts observed in our results agrees with previous studies, particularly [19], who also found that species were affected more severely as the shifts grow larger. That study reported that, whenever species phenophases are estimated in the same way we did, phenological shifts of 1 to 3 weeks are predicted to cause ∼ 17 − 30% of pollinator species being affected in their interactions with plants. Our results qualitatively agree with these predictions, although we explored a wider range of phenological shifts: between −6 to 6 weeks.

From our results we could also conclude that evaluating shifts in only mean phenological values, i.e. without considering the phenological variance, can be insufficient to understand the effects of phenological disruptions. This comes from observing that changing the inter-annual variability in the magnitudes of phenological shifts strongly affects species persistence, even though the mean value of the shifts over all species was zero. A similar result was previously reported from experimental manipulations of life-history events in competing tadpoles [38].

Regarding the underlying causes of species persistence, our analysis revealed that phenophase duration in plants, along with both phenophase duration and degree of pollinators, are key attributes determining species persistence in the long run. In agreement with our results, Fagan *et al.* [21] found that longer flowering phenophases favored plant persistence, while Memmott *et al.* [19] found that higher connectivity of pollinators led to higher pollinator persistence. Our analysis replicated both previous results with a single modeling approach.

In the datasets used in our study, species degree and the length of their phenophases are positively correlated (see S3 Table). However, these correlations are stronger for pollinators than for plants. In the context of our model, we can provide an explanation for the importance of phenophase duration and why degree is specially important for pollinators to overcame phenological shifts. Phenophase length may increase persistence probability of species because a longer phenophase implies more opportunities for finding an interacting species at any given time. In a context of phenological shifts, longer phenophases decrease probability of losing interactions. However, plants and pollinators exhibit some important differences in the functional significance of phenophase length. Plants with longer phenophases tend to produce more flowers and rewards for pollinators, promoting both seed and larvae production. In addition, for plants there are no trade-offs between phenophase length and reproduction success. This is because plants can sustain flower abundance through time (see first term of Eqn. (4)). In contrast, if pollinators lengthen their activity period reproductive rate may drop due to decreasing abundance of reproductive adults (Eqn. (8)). For this reason, to maintain larval production, pollinators need to combine high connectivity with long phenophases to compensate for decreases in abundance. This can explain the strong correlation observed between degree and phenophase length in pollinators.

Note that degree and abundance are usually directly related in both plants and pollinator species in a given network [39]. Furthermore, the presence of low–abundance species tends to be underestimated since detecting them requires a greater sampling effort [40]. Consequently, the degree and phenophase duration of specialist (i.e. rare) species are likely to be underestimated [40–42]. Based on our findings, this imply that ultra–specialist species are more scarce and therefore their persistence probability could be higher than suggested by recorded data.

In this study we have considered a rather wide gradient of phenological shifts for our analyses. However, empirical estimations show an average advance of 11.5 days in 120 years (roughly 1 day per decade) for appearance of pollinators, and 9.5 days for plant flowering in America [13]. Likewise, estimates for European pollinators suggest an average phenological advancement of 6 days in 60 years (1 day per decade) [43]. Such amounts of phenological shifts are consistent and might imply a rather small current degree of temporal mismatch of interactions, with low rates of predicted extinctions based on our results. Nevertheless, if phenological shifts accelerate as a consequence of climate changes and cross the threshold of 2 weeks at the community level, our results predict an abrupt increase in extinction rates.

Current empirical evidence supports that the timing of flowering and pollinator emergence are being disrupted, presumably due to climate changes [7, 44]. These observed shifts in phenological events could lead to temporal mismatches between flowering seasons of species and the periods of pollinator activity (as suggested in [8]), although direct empirical evidence of this kind of interaction disruption is elusive up to date [45, 46]. For example, [31] and [47] showed that visitation frequency, associated to pollinator abundances, decreased along with observed flowering advancements. Likewise, [48] showed that early flowering produced lower pollination success and consequent lower reproductive success for plants. Schenk *et al.* [49] found that even small mismatches between plants and pollinators affect reproduction, activity and survival of pollinators. However, the consequences of phenological shifts and the interaction disruptions remains to be understood at the community and ecosystem levels. Our study contributes to filling a gap in the understanding of community level effects of shifting phenologies in plant–pollinator interactions [50]. However, although we identified consistent responses from the analysis of some perturbed plant–pollinator systems, our results show considerable differences among the studied networks. This underlines the fact that network studies based on a single community should be interpreted with caution since revealing general patterns requires a comparative analysis among networks. Our findings, in agreement to observed in previous work [51], highlight the importance of phenologies on the stability and robustness of mutualistic communities. Future research should consider studying some real phenomena not included in our model. For example, the shortening of the pollination season already observed in some real life systems [43, 52]. Additionally, between-community variability in responses to phenological shifts poses the questions about which are the chief determinants of these differences, species traits or connectivity patterns.

## Supporting information

**S1 Table. Initial and boundary conditions of state variables.** For each species, we divided the shown initial value by the number of species in its group.

**S2 Table. Model parameters.** Symbols, short description and mean values.

**S3 Table. Correlation between topological degree and phenophase duration.** Pearson and Spearman correlation values between topological degree and phenophase duration for plants and pollinator for every network. All values are statistically significant (*P* < 0.05) except Spearman’s correlation for plant species in Nahuel Huapi (in italics).

**S1 Fig. NMDS analysis of persistence differences among networks.** Results of non-metric multidimensional scaling (NMDS) analysis on persistence for the 8 networks under study, belonging to three locations: Zackenberg (blue), Nahuel Huapi (red), and Villavicencio (green).

**S2 Fig. NMDS analysis of importance differences among networks.** Results of non-metric multidimensional scaling (NMDS) analysis on importance attribute for the 8 networks under study, belonging to three locations: Zackenberg (blue), Nahuel Huapi (red), and Villavicencio (green).

**S1 Appendix. Plant and pollinator species persistence under different modes and magnitudes of phenological shift.**

**S2 Appendix. Importance measures of species attributes for the long–term persistence of pollinators and plants.** Results are presented for the different locations. The panels show different phenological change scenarios, modeled by different temporal shifts distribution, *TSD* (see Methods): *TSD–msd* (B), *TSD–m* (M), *TSD–sd* (V), *TSD–msdy* (Y). Magnitude of phenological shifts are presented in the x—axes.

## Acknowledgments

The authors thank Jens Olesen for providing the Zackenberg dataset used in our analyses.

